# Far-red and sensitive sensor for monitoring real time H_2_O_2_ dynamics with subcellular resolution and in multi-parametric imaging applications

**DOI:** 10.1101/2024.02.06.579232

**Authors:** Justin Daho Lee, Amanda Nguyen, Zheyu Ruby Jin, Aida Moghadasi, Chelsea E. Gibbs, Sarah J. Wait, Kira M. Evitts, Anthony Asencio, Samantha B Bremner, Shani Zuniga, Vedant Chavan, Andy Williams, Netta Smith, Michael Regnier, Jessica E. Young, David Mack, Elizabeth Nance, Patrick M. Boyle, Andre Berndt

**Affiliations:** Molecular Engineering and Sciences Institute, University of Washington, Seattle, WA, USA; Department of Bioengineering, University of Washington, Seattle, WA, USA; Institute for Stem Cell and Regenerative Medicine, University of Washington, Seattle, WA, USA; Department of Laboratory Medicine and Pathology, University of Washington, Seattle, WA, USA; Department of Chemical Engineering, University of Washington, Seattle WA, USA; Center for Cardiovascular Biology, University of Washington, Seattle WA, USA; Department of Rehabilitation Medicine, University of Washington, Seattle, WA, USA

## Abstract

H_2_O_2_ is a key oxidant in mammalian biology and a pleiotropic signaling molecule at the physiological level, and its excessive accumulation in conjunction with decreased cellular reduction capacity is often found to be a common pathological marker. Here, we present a red fluorescent Genetically Encoded H_2_O_2_ Indicator (GEHI) allowing versatile optogenetic dissection of redox biology. Our new GEHI, oROS-HT, is a chemigenetic sensor utilizing a HaloTag and Janelia Fluor (JF) rhodamine dye as fluorescent reporters. We developed oROS-HT through a structure-guided approach aided by classic protein structures and recent protein structure prediction tools. Optimized with JF_635_, oROS-HT is a sensor with 635 nm excitation and 650 nm emission peaks, allowing it to retain its brightness while monitoring intracellular H_2_O_2_ dynamics. Furthermore, it enables multi-color imaging in combination with blue-green fluorescent sensors for orthogonal analytes and low auto-fluorescence interference in biological tissues. Other advantages of oROS-HT over alternative GEHIs are its fast kinetics, oxygen-independent maturation, low pH sensitivity, lack of photo-artifact, and lack of intracellular aggregation. Here, we demonstrated efficient subcellular targeting and how oROS-HT can map inter and intracellular H_2_O_2_ diffusion at subcellular resolution. Lastly, we used oROS-HT with the green fluorescent calcium indicator Fluo-4 to investigate the transient effect of the anti-inflammatory agent auranofin on cellular redox physiology and calcium levels via multi-parametric, dual-color imaging.

## Introduction

Oxidative stress is often a key component of many disease progressions. Tremendous efforts have been made to develop therapeutic approaches to target the excessive presence of oxidants and their source. However, the unsatisfying results of antioxidative therapy call for a more nuanced understanding of cellular oxidants, antioxidative defense networks, and their effect on the cellular system with precision and specificity to improve rationales on antioxidative therapeutics^1^.

H_2_O_2_ is a major oxidant in redox biology that can also act as a pleiotropic secondary messenger in various cellular signaling processes^2–6^. Its precursor superoxide is a natural byproduct of aerobic metabolism, which rapidly gets converted to H_2_O_2_ naturally or by superoxide dismutases (SOD)^7^. The level of intracellular H_2_O_2_ is tightly regulated by peroxide-reducing mechanisms. Although peroxide is considered less reactive than other cellular oxidative agents, its excessive accumulation is often observed in pathology, with growing evidence of its causal role in the progression of diseases^8–10^. The engineering of genetically encoded H_2_O_2_ indicators (GEHI, e.g. HyPer^11,12^, roGFP family sensors^13,14^) has been a significant step towards understanding the role of peroxide in redox biology by enabling real-time monitoring of peroxide dynamics in a wide array of biological hosts^15^. One advantage of GEHIs over redox-sensitive fluorescence dyes is their spatiotemporal flexibility: they can be targeted to specific cell types or various cellular compartments for extended periods when coupled with proper expression systems (e.g. promoters and trafficking/export tags). Specifically, red-fluorescent GEHIs facilitate multiparametric analysis of peroxide dynamics along with other key biomolecules or processes considering a large number of green fluorescent sensors for biological molecules and processes (e.g. Ca^2+^, pH, voltage, redox potential, etc.)^16,17^. Nevertheless, current red-shifted GEHIs exhibit slow kinetics, a bottleneck for real-time peroxide imaging. Most importantly, current red GEHIs exhibit a blue-light-induced photochromic artifact, making unobstructed multiparametric analysis alongside green fluorescent sensors difficult. Lastly, aggregation tendency and low brightness are also common among red fluorescent proteins^18^ and affect the performance of existing red GEHIs.

In this study, we coupled the bacterial OxyR peroxide sensor with a rhodamine-HaloTag-based chemigenetic reporter system to create a first-in-class, far-red indicator for H_2_O_2_: oROS-HT_635_ (optogenetic hydRogen perOxide Sensor with HaloTag with JF635). We developed a rational engineering strategy based on structural information derived from experimentally resolved structures and computational methods (ColabFold)^19^. oROS-HT_635_ has excitation and emission wavelengths of 640 nm and 650 nm. We validated it in various biological host systems, including stem cell-derived cardiomyocytes *in vitro* and primary neurons *ex vivo*. Moreover, we found that the fast oROS-HT_635_ kinetics allows the observation of intracellular diffusion of peroxide. Also, oROS-HT_635_ is free from photochromic false positive artifacts, allowing multiparametric analysis of contextual peroxide dynamics. As a proof-of-concept, we showed the acute effect of the anti-inflammatory agent auranofin on peroxide with the context of changes in cellular redox potential in HEK293 cells and Ca^2+^ in human induced pluripotent stem cell-derived cardiomyocytes (hiPSC-CMs), demonstrating the intriguing effect of antioxidant system perturbation in acute peroxide increase.

## Results

### Structure-guided engineering of oROS-HT_635_: a bright far-red optogenetic sensor for H_2_O_2_

OxyR is a bacterial transcription activator with high specificity and sensitivity toward H_2_O_2_ with low peroxidative capability (i.e. the protein exhibits high sensitivity towards peroxide with limited catalytic activity)^20^. Existing red-shifted GEHI, such as HyPerRed^12^ and SHIRMP^21^ utilize ecOxyR-LBD (regulatory domain of OxyR from Escherichia coli), as their sensing domain. However, both red GEHIs show slower kinetics (10s to 100s seconds for full activation under saturation and half an hour for reduction) than the innate kinetics and sensitivity reported for ecOxyR itself^22–24^. Specifically, ecOxyR oxidation of ecOxyR is at a sub-second scale, and its reduction takes 5∼10 minutes, implying that the insertion of the fluorescence reporter domain slowed down the activation and deactivation of ecOxyR. Our engineering strategy aimed to maintain the flexibility of the protein loop that drives the conformational change in ecOxyR in the derived sensors as we previously described for a GFP-based oROS sensor^25^. Specifically, ecOxyR contains a hydrophobic pocket that forms the active center for peroxide interactions. Upon binding, peroxide forms a hydrogen bonding network with adjacent residues, bringing residues C199 and C208 into close proximity to form a disulfide bridge. By analyzing the B-factors of OxyR structures, we observed high flexibility in the 205-222 region of ecOxyR **[Fig. 1A]**. We reasoned that preserving this flexibility is necessary for efficient OxyR activation by peroxide. Thus, inserting a bulky fluorescent reporter between C199 and C208, as in HyPerRed and SHIRMP, may significantly slow OxyR’s activation, and we tested alternatives outside this region **[Fig. 1B]**. Furthermore, red fluorescent proteins pose challenges for versatile use involving optical multiparametric analysis or neuron expression. For example, cpmApple, used in HyPerRed, exhibits a false positive photochromic artifact induced by blue light commonly used to excite green fluorescent proteins (e.g. 488 nm).

**Figure 1.**
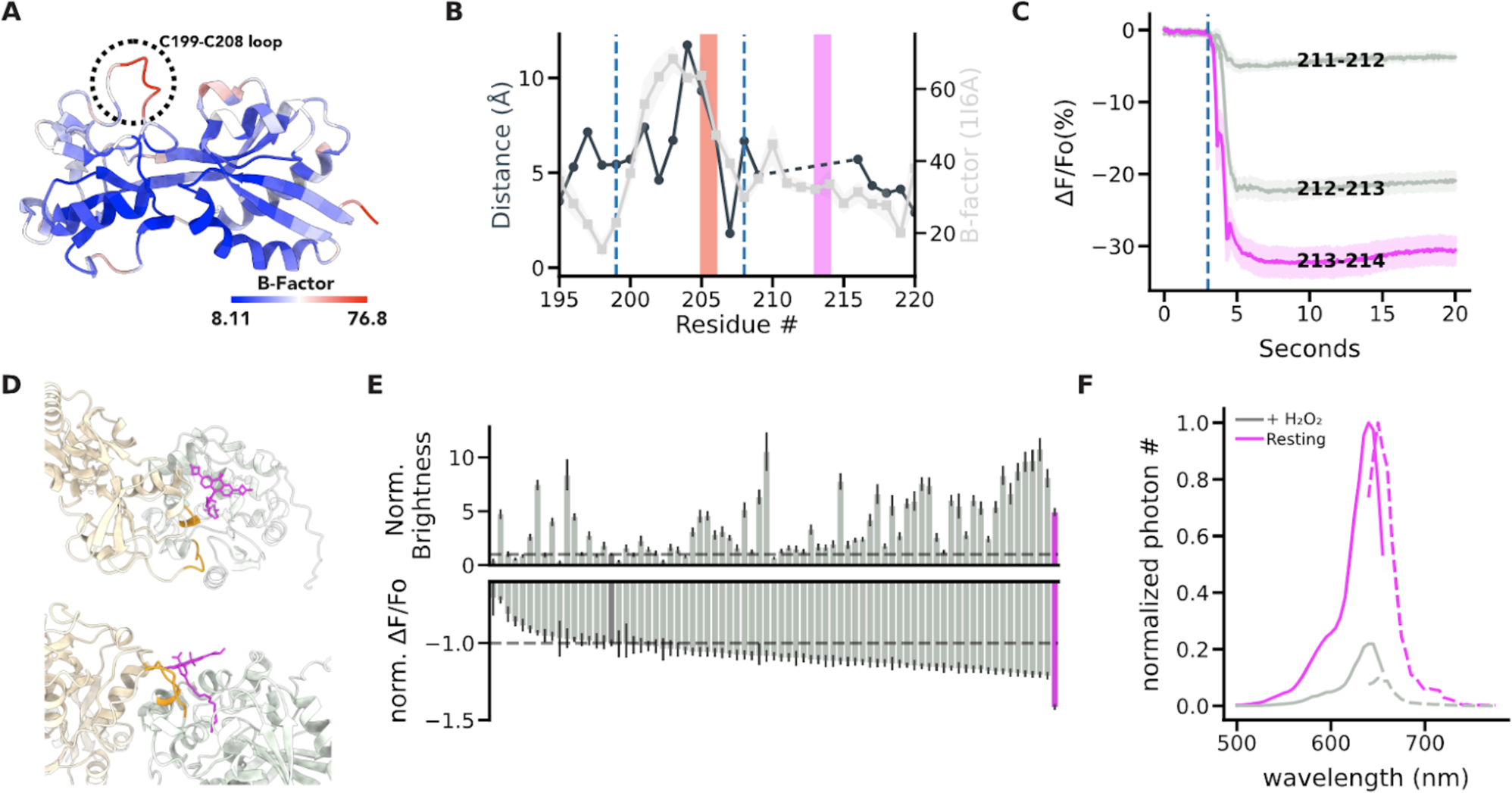
Structure-guided engineering of oROS-HT: bright far-red optogenetic sensor for H_2_O_2_. **A** B-Factor of the regulatory domain (RD) of the oxidized form of E. coli OxyR [PDB:1I6A] indicates flexible and rigid regions. The flexible loop between C199-C208 facilitates disulfide bridging of C199 and C208 residues. **B** Pairwise distance between residues #195 to #220 of oxidized [PDB:1I6A] and reduced [PDB:1I69] crystal structures of ecOxyR RD (Gray line). Average B-factor of residues #195 to #220 in oxidized ecOxyR RD [PDB:1I6A] (Black line). residues #195 to #220 indicate a putative region with high conformational change in ecOxyR. The dotted lines indicate C199 and C208. The red and pink bars indicate the insertion sites of fluorescence reporters in HyPerRed (cpmApple) and oROS-HT (cpHaloTag), respectively. **C** Prototype variants of oROS-HT from insertional screening between #210 to # 215, a novel putative site for reporter insertion in ecOxyR. Fluorescence change (ΔF/Fo %) in response to extracellular H_2_O_2_ (300µM) stimulation on the variants expressed in HEK 293 WT cells. (Cell of interests collected from 3 biological replicates. 211-212 [n = 248], 212-213 [n= 386], 213-214 [n = 249]) **D** Putative JF635 ligand position bound to predicted oROS-HT structure in 2 different perspectives. ColabFold predicted structure of the 213-214 cpHaloTag variant was superimposed with cpHaloTag (PDB:6U2M) crystal structure to obtain a relative position of JF635 on the 213-214 cpHaloTag variant. *gray*: cpHaloTag (reporter domain) *beige*: OxyR (sensing domain) *orange*: linkers, *magenta:* JF635-HTL. **E** Random linker mutagenesis screening of linkers between both domains (X-cpHaloTag-XX) in HEK 293 WT cells. ***top*** Normalized brightness of each sensor variant. ***bottom*** Normalized ΔF/Fo (213-214 variant at −1.0) of each sensor variant with 300µM H_2_O_2_ stimulation. (211-212 [n=1468], oROS-HT [n=1218], the rest of the sample numbers can be found in the supplementary table 1. Magenta = oROS-HT, Dark Gray = 211-212, and dashed lines = mean of 211-212 **F** Spectral profile of oROS-HT expressed in HEK293. Solid line: excitation spectra (peak: 640nm), dotted line: emission spectra (peak: 650nm). **Statistics:** Error bars and bands represent the bootstrap confidence interval (95%) of the central tendency of values using the Seaborn (0.11.2) statistical plotting package.

Deo et al. proposed a chemigenetic solution incorporating a self-labeling enzyme (HaloTag) labeled with rhodamine-based red-shifted Janelia Fluorophores (JF)^26,27^. JFs were designed for wavelengths between 494nm and 722nm and exhibit exceptional photophysical characteristics such as brightness, and photostability, and low pH sensitivity. We aimed to engineer a new class of GEHIs using cpHaloTag labeled with the red fluorescent JF635 as a reporter. Insertion of cpHaloTag into multiple positions outside of the C199-C208 loop in ecOxyR was well tolerated, and we identified a prototype sensor variant 213-214 with a robust response to extracellularly introduced 300µM H_2_O_2_ (ΔF/Fo%: −38.23%; ci = [−40.36, −36.18]) **[Fig. 1C]**. Interestingly, we observed inverse responses (e.g. increase in peroxide level leads to decreased fluorescence) to peroxide in all insertional variants. Thus, we aimed to improve the resting brightness, guided by using structure prediction from ColabFold (AlphaFold2 with MMseqs2 for multiple sequence alignment). The prediction yielded a highly confident structure of variant 213-214, which is exemplified by a dimeric interface of the sensing domain that closely resembles the dimeric interface of reduced ecOxyR resolved by crystallography **[Supp. Fig. 1A, B].** We superimposed the cpHaloTag-JF635 structure from [PDB: 6U2M] to identify the putative position of JF635 with the sensing domain of variant 213-214 **[Supp. Fig. 1C].** We found the OxyR sensing domain in a position oriented away from JF635 rather than covering the JF635 fluorophore **[Fig. 1D]**, increasing the putative influence of interdomain linker regions as the fluorophore’s local environment. This configuration is consistent with the spatial configuration of the chemigenetic calcium indicator HaloCaMP^26^.

Consequently, random mutagenesis of interdomain linker residues (XX-cpHaloTag-X, X indicates mutagenesis targets) affected both the sensor brightness and dynamic range **[Fig. 1E]**. From the linker variant library, we found a variant with 4.9-fold increased resting brightness and a 41% increase in dynamic range induced by 300µM H_2_O_2_ compared to those of variant 213-214 (resting brightness: relative fluorescence intensity, variant 213-214: 160.71; ci = [153.03, 168.79], oROS-HT_635_: 788.24; ci = [742.26, 834.7]; dynamic range: 213-214: 160.71 (n = 1468); ci = [153.03, 168.79], oROS-HT_635_: −67.99 (n = 1218); ci = [−68.52, −67.45]), which was later named oROS-HT_635_ **[Fig. 1F]**. In addition to the structural hypothesis of the interdomain linker’s influence on both sensor dynamics and brightness, we also identified F209 to be a putative mutational site for the fluorophore local environment tuning, resulting in a more than a 3-fold difference in resting brightness between the dimmest variant (F209L) and the brightest variant (F209R) and the trend was also consistent when the sensor was labeled with ligand JF585. Unfortunately, the mutational benefit of MS-cpHaloTag-N and F209R was non-synergistic, which led us to exclude mutation F209R for our final variant **[Supp. Fig. 1D-G]**.

### Characterization of ultrasensitive and fast H_2_O_2_ sensor, oROS-HT**_635_**

We first characterized oROS-HT_635_ by exogenously applying H_2_O_2_ to cells expressing the sensor and second by applying menadione, which induces intracellular peroxide generation. Menadione generates H_2_O_2_ through various redox cycling mechanisms^28–31^ **[Fig. 2A]**. Saturation of oROS-HT_635_ induced by 300µM H_2_O_2_ revealed a fast sub-second activation that could capture the H_2_O_2_ diffusion across the imaging field of view. It implies that the kinetic efficiency of the sensor passed a milestone of no longer being reaction-limited in this scenario. Intriguingly, the response amplitude of oROS-HT_635_ at 10µM external peroxide was −58.69% ΔF/Fo (ci = [−59.18, −58.18]), which is 87% of the amplitude at saturation upon 300µM peroxide (−67.27% ΔF/Fo; ci = [−67.64, −66.91])), demonstrating the high sensitivity of the sensor **[Fig. 2B]**. Previous studies showed the intracellular H_2_O_2_ concentrations in HEK293 cells are at approximately 10 and 300 nM under these external conditions, respectively^12,32^. Furthermore, oROS-HT_635_ allowed the monitoring of titrated peroxide levels in HEK293 cells induced by 10, 20, and 50µM of menadione. We measured a concentration-dependent response in oROS-HT_635_ signal of −26.8% ΔF/Fo (ci = [−27.63, −25.98]), −59.59% ΔF/Fo (ci = [−60.48, −58.67]), and −63.06% ΔF/Fo (ci = [−63.59, −62.51]) in ΔF/Fo, respectively **[Figure. 2C]**. Interestingly, under 50 µM menadione, oROS-HT_635_ reaches near maximum fluorescence amplitudes but at much slower rates than exogenously induced instant H_2_O_2_ saturation (300 uM). Therefore, these kinetics most likely show the real-time increase of cytosolic peroxide by menadione.

**Figure 2.**
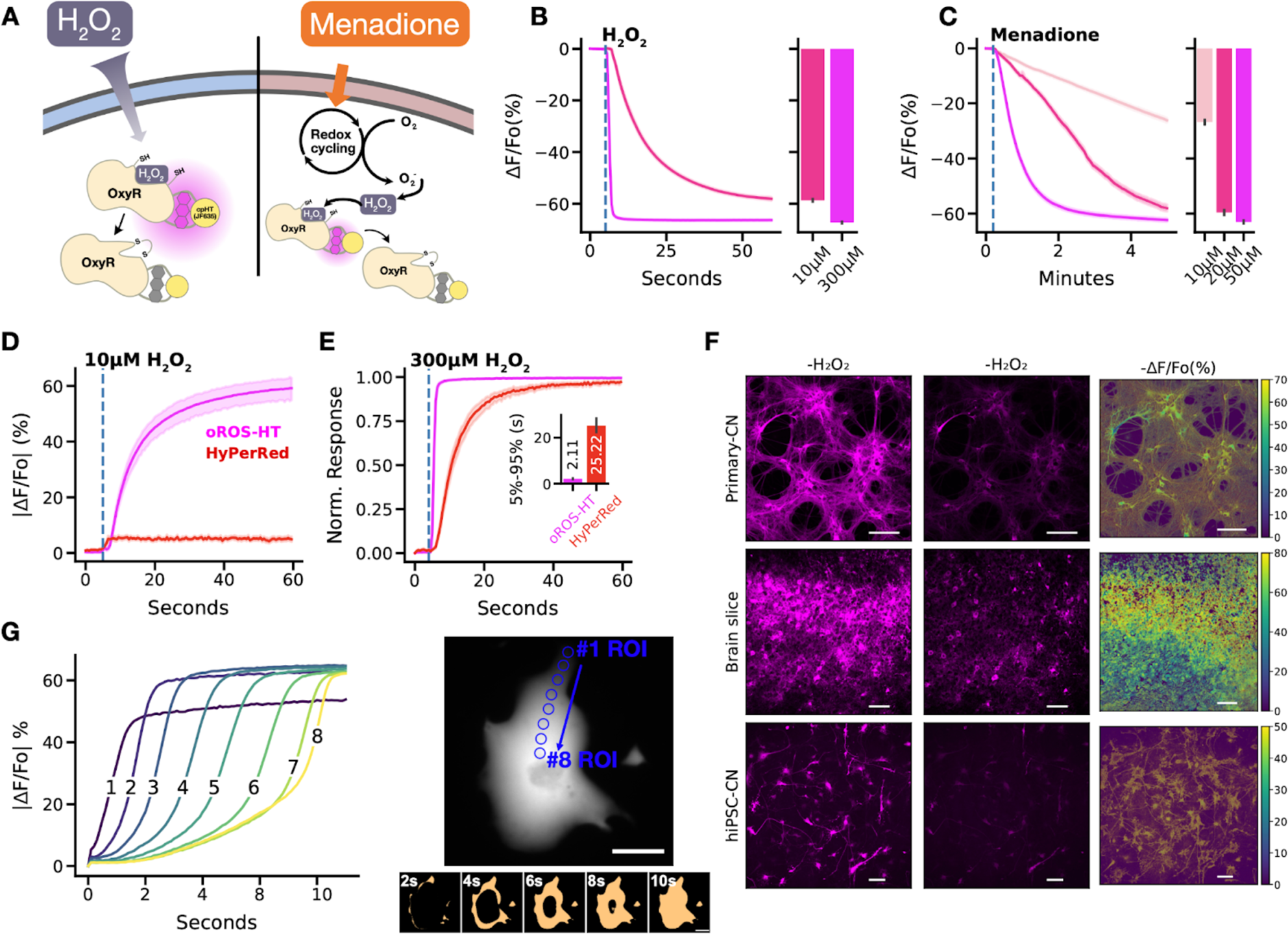
Characterization of ultrasensitive and fast H_2_O_2_ sensor, oROS-HT. **A-C** oROS-HT sensor characterization. **A** Schematic illustration of the methods. oROS-HT’s fluorescence change was characterized by its response to either exogenous H_2_O_2_ shot (exogenous H_2_O_2_ Figure 2B) or Menadione (cell-sourced H_2_O_2_ Figure 2C). Menadione goes through an NADPH oxidase-like redox cycle to produce intracellular H_2_O_2_ **B** Fluorescence changes of oROS-HT upon exogenous administration of high (300µM) or low (10µM) H_2_O_2_. oROS-HT was expressed in HEK293 WT cells (n>100 cells each). The barplot represents the mean of the maximum fluorescent response of ROI. **C** Fluorescence response of oROS-HT in HEK293 to varying concentrations of menadione (n>100 cells each). The barplot represents the mean of the maximum fluorescent response of ROIs. **D-E** Benchmarking oROS-HT with existing red H_2_O_2_ sensor HyPerRed. **D** Representative response of oROS-HT and HyPerRed to 10µM exogenous H_2_O_2_ (n=31, cells per sensor) **E** Normalized response of oROS-HT and HyPerRed to exogenous 300µM H_2_O_2_ (n=23, cells per sensor). **F** Expression of oROS-HT in primary cortical neurons (Primary-CN), *ex vivo* rat cortex (Brain slice), and hiPSC-derived cortical neurons (hiPSC-CN) and their responses exogenous 300µM H_2_O_2_. Scale bars: 100µm **G** Intracellular diffusion of H_2_O_2_ across an iPSC-CM expressing oROS-HT captured at 20Hz frame rate upon 300µM H_2_O_2_ extracellular administration. ***right*** Pixel values are transformed to “False” at 0∼50% sensor activation and “True” at 50%∼100% sensor activation to enhance visualization. Scale bar = 50µm ***left*** Absolute sensor response magnitude and kinetics at 8 ROIs within the cardiomyocyte during the intracellular diffusion of H_2_O_2_. Blue circles on the right image indicate the ROIs used for the plots on the left. **Statistics:** Error bars and bands represent the bootstrap confidence interval (95%) of the central tendency of values using the Seaborn (0.11.2) statistical plotting package. Cell-of-interests were collected from 3 biological replicates unless noted otherwise.

Next, we conducted a benchmark validation in HEK293 cells with the current best-in-class red GEHI HyPerRed (ex/em 575/605nm) to illustrate the improvements made in oROS-HT_635_. Despite being an inverse response sensor, the magnitude of absolute fluorescence change from the baseline of oROS-HT_635_ in response to low-level (10µM) peroxide stimulation was **≈**59%, where the same condition only caused below 5% change in HyPerRed **[Fig. 2D]**. Additionally, the oROS-HT_635_ response kinetics were approximately 8 times faster under saturating 300 uM peroxide compared to HyPerRed (5-95% |ΔF/Fo| time, oROS-HT_635_: 0.96 s; ci = [0.87, 1.04], HyPerRed: 7.8 s; ci = [6.98, 8.72]) **[Fig. 2E]**. oROS-HT_635_ also displayed robust expression in various mammalian tissues (e.g. primary rat cortical neurons and *ex vivo* rat brain tissue) and human stem cell-derived models (e.g. cardiomyocytes and cortical neurons) **[Fig. 2F]**. Furthermore, cytosolic oROS-HT_635_ revealed peroxide diffusion at **≈**10µm/s in cardiomyocytes when exposed to 300µM H_2_O_2_ **[Fig. 2G]**. For the first time, we optically monitored the influx of H_2_O_2_ into hiPSC-CMs with subcellular resolution, demonstrating that the sensor dynamics reflect the diffusion event. Many experimental studies of intracellular peroxide often assume well-mixed uniformity of peroxide concentrations^15,32^. However, a previous model for cytosolic H_2_O_2_ also showed spatial peroxide gradients in mammalian cells can emerge upon external peroxide stimulation^32^ consistent with our observation.

### Optimized biophysical properties and versatility of oROS-HT_635_ under varying conditions

We envision users of oROS-HT_635_ studying peroxide dynamics under varying conditions. Thus, we further characterized notable features of oROS-HT_635_ that demonstrate its environmental resiliency. oROS-HT_635_ could be repeatedly activated and reduced back to baseline by serial peroxide stimulation and washout, demonstrating the reversibility of the sensor. Thus, the sensor is able to track real-time fluctuations of intracellular peroxide **[Fig. 3A, B]**. Most beta-barrel fluorescent proteins in sensor designs require oxygen for their fluorophore maturation^33,34^. In addition, it was reported that GFP undergoes photoconversion under hypoxic conditions, where the excitation/emission spectra shift and become similar to RFP^35^. In contrast, the HaloTag-Rhodamine-based chemigenetic sensors utilize a synthetic fluorophore, making their maturation independent from oxygen levels. To demonstrate oxygen independence during maturation, we engineered a loss-of-function mutation of oROS-HT_635_ (C199S), a sensor variant insensitive to peroxide **[Fig. 3C]**. As a negative control, oROS-HT_635_-C199S can reflect any environmental effect on the level of fluorescence that is not associated with the sensor function^12^. HEK293 cells transfected with oROS-HT_635_-C199S did not significantly differ in expression level when matured under normoxic or hypoxic conditions **[Fig. 3D-F]**. Red-shifted GEHIs are often limited for multiparametric use with green sensors due to a photochromic false positive artifact in response to blue light. However, oROS-HT_635_ lacks this artifact, rendering oROS-HT_635_ ideally compatible with green reporters **[Fig. 3G]**. Harnessing its multiplexing capability, we co-expressed oROS-HT_635_ or oROS-HT_635_-C199S with a GFP-based pH indicator SypHer3s to demonstrate the low pH sensitivity of oROS-HT and its functionality under pH change with sequential events of 1.) acidic pH insult (pH 6) and 2.) 10µM menadione-induced peroxide increase. oROS-HT_635_ did not respond to the initial change in pH but detected the menadione-induced increase in cytosolic peroxide, exemplifying its robust functionality under changing cellular pH environments **[Fig. 3H]**. As a benchmarking comparison, we compared pH-dependent fluorescence change of oROS-HT-C199S and HyPerRed-C199S under neutral pH (pH 7.44) in response to pH shift to either 9 (basic) or 6 (acidic). oROS-HT_635_-C199S exhibited no significant fluorescence change to either condition compared to HyPerRed-C199S at equivalent conditions, demonstrating that the oROS-HT’s fluorescence is largely insensitive to physiological pH fluctuation in contrast to HyPerRed **[Fig. 3I, J]**.

**Figure 3.**
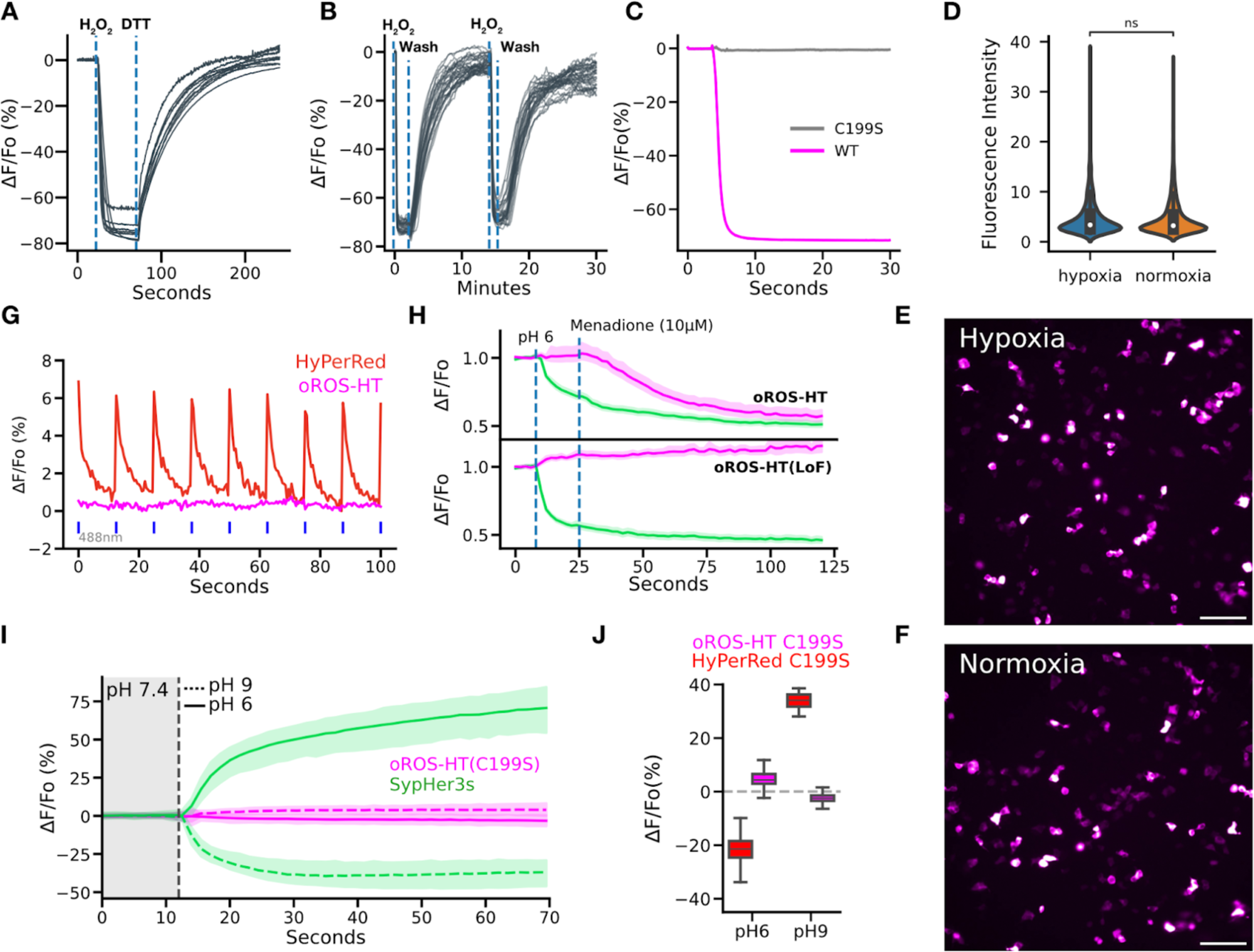
Optimized biophysical properties and versatility of oROS-HT_635_ under varying external conditions. **A-B** Reversibility of oROS-HT **A** HEK293 expressing oROS-HT_635_ was first stimulated with 100µM H_2_O_2_ and then 10 mM Dithiothreitol (DTT), a reducing agent, shortly after. **B** HEK293 expressing oROS-HT were stimulated with 100µM H_2_O_2_ followed by media wash and 2nd stimulation. (n=32, cells) **C** Fluorescence change of oROS-HT_635_ is H_2_O_2_ specific. Fluorescence response of oROS-HT_635_-WT and oROS-HT_635_-C199S expressed in HEK293 to 300µM H_2_O_2_. C199S mutation disables OxyR’s H_2_O_2_ specific C199-C208 disulfide bonding mechanism. oROS-HT_635_-C199S can be utilized as a negative control sensor. **D-F** Maturation of oROS-HT(C199S) in a hypoxic condition. HEK293 expressing Loss-of-function oROS-HT(C199S) were incubated for 18 hours in either Normoxia (atmospheric condition at 37°C) or Hypoxia (N2 infused chamber at 37°C) overnight (18h). **D** Fluorescence intensity profile of oROS-HT(C199S) (Hypoxia [n = 1246] / Normoxia [n=1765] collected from 8/11 biological replicates, respectively). **E,F** Representative images oROS-HT(C199S) maturated in HEK293 cells in Hypoxia or Normoxia conditions. Scale bar = 100µm. **G** Representative fluorescence emission of oROS-HT and HyPerRed under their respective excitation wavelength (635nm and 597nm) and in response to 488nm light pulses. **H** Dual monitoring of pH and H_2_O_2_ in mammalian cells. Either oROS-HT or oROS-HT(C199S) were paired with SypHer3s, a green fluorescence pH indicator, to be co-transfected on HEK 293 cells to monitor the sequential events of 1. pH environment change (pH6) and 2. Menadione (10µM) induced H_2_O_2_ increase. (n>100 per condition)**. I** Multiplexed epifluorescence imaging of Loss-of-function oROS-HT(C199S) and SypHer3s coexpressed in HEK293 cells. Neutral imaging solution (PBS, pH 7.44) was switched to either acidic (PBS, pH6) or basic (PBS, pH9) imaging solution at the vertical dashed line (gray). **J** ΔF/Fo (%) of oROS-HT(C199S, from Fig. 3I) and HyPerRed(C199S) at pH9 or pH6. Left and right box plots for each condition represent values at the first and last frames, respectively. **Descriptive Statistics:** Error bars and bands represent the bootstrap confidence interval (95%) of the central tendency of values using the Seaborn (0.11.2) statistical plotting package. F: Error bands represent the 95% interval, ranging from the 2.5 to the 97.5 percentiles from medians using the Seaborn (0.12.1) statistical plotting package. Cell of interest collected from 3 biological replicates. SypHer3s pH9: 59.56 % ΔF/Fo (n = 883); ci = [58.46, 60.62]. SypHer3s pH6: −34.56 % ΔF/Fo (n >100); ci = [−35.15, −33.96]. oROS-HT(C199S) pH9: −2.43 (n >100); ci = [−2.54, −2.32]. oROS-HT(C199S) pH6: 4.9 (n > 100); ci = [4.64, 5.19]. **Inferential Statistics:** D: t-test independent samples. *P < 0.05, **P < 0.01, ***P < 0.001.

### Multiparametric analysis of the acute effect of auranofin on H_2_O_2_, redox potential, and Ca^2+^

#### Acute effect of auranofin on cellular H_2_O_2_ level and redox potential

Grx1-roGFP2 is an indicator sensitive to glutathione redox potential (E_GSH_). It is a fusion between glutaredoxin1 (grx1) and the redox-sensitive green fluorescent protein roGFP2. Multiplexed imaging of oROS-HT_635_ with Grx1-roGFP2 could enable peroxide imaging with augmented information about the redox cellular environment. Here, we monitored both sensors simultaneously in HEK293 cells upon 10µM H_2_O_2_ exposure. We revealed sequential events of intracellular peroxide increase followed by a decrease in glutathione redox potential E_GSH_ (peak_oROS-HT_ to peak_Grx_ = 3.12 s) as indicated by the respective sensor responses **[Fig. 4A]**. In contrast, inhibition of cellular redox potential with Trx/Grx (Thioredoxin/Glutaredoxin) inhibitor auranofin (1µM) showed rapid decay of E_GSH_ followed by a slow increase of intracellular peroxide level. Interestingly, auranofin-induced peroxide build-up was transient, as we observed the elevation in peroxide level for 45 minutes after the application, followed by a recovery to the baseline within the following 60 min **[Fig. 4B],** potentially due to stress-induced antioxidative capacity increase. Consistent with the previous reports^36,37^, we observed increased translocation of Nrf2 into the nucleus within 30 minutes of exposure to 1µM Auranofin **[Fig. 4C, Supp. Fig. 2].** In conclusion, the multiplexed use of Grx1-roGFP2 with oROS-HT_635_ exemplifies the peroxide monitoring capability of oROS-HT_635_ in the context of the cellular redox environment.

**Figure 4.**
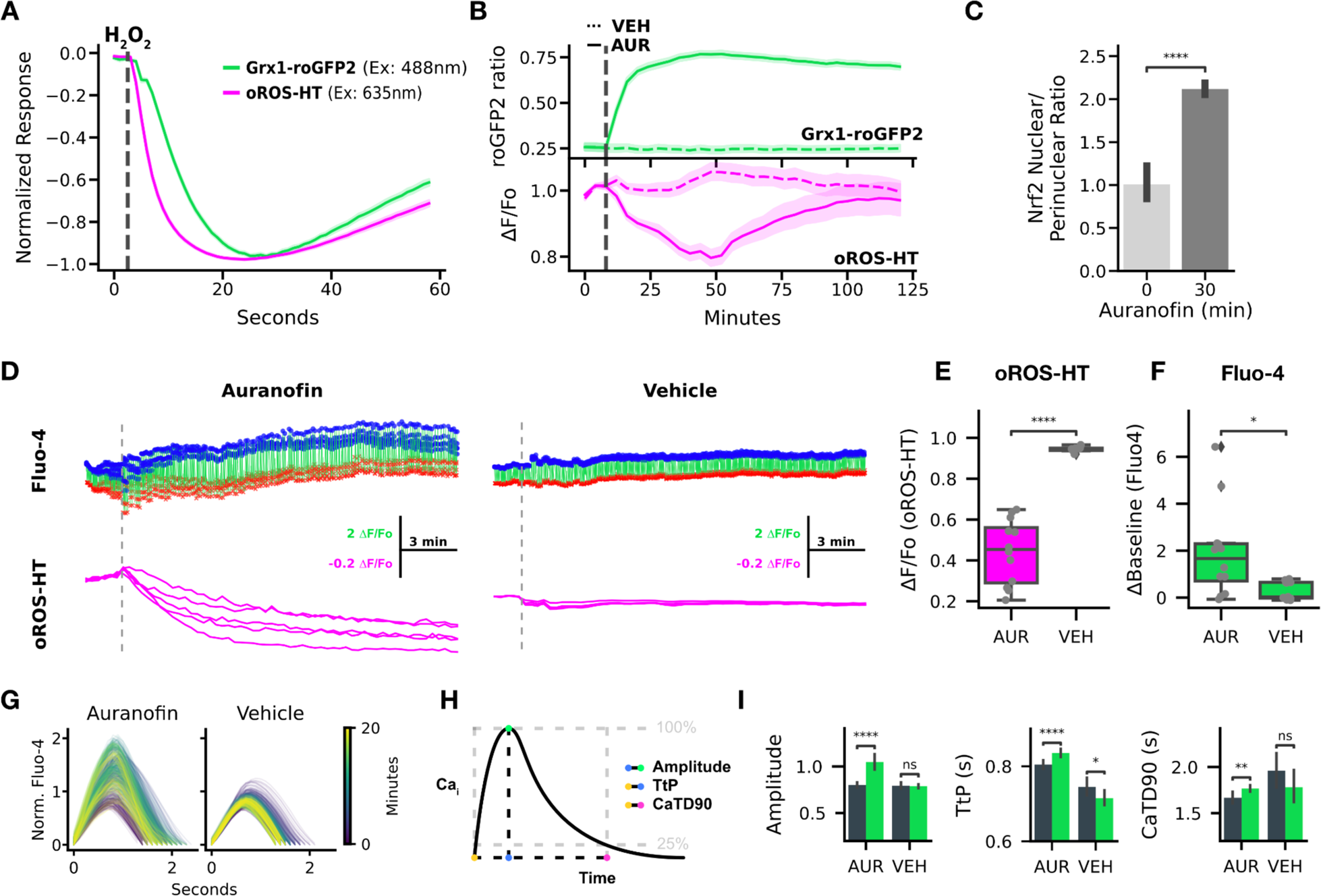
Multiparametric analysis of the acute effect of auranofin on H_2_O_2_, redox potential, and Ca^2+^. **A-C** Dual monitoring of intracellular glutathione redox potential and H_2_O_2._ **A** Normalized fluorescence change of Grx1-roGFP2 (green, glutathione redox potential sensor) and oROS-HT (magenta) co-expressed in HEK293 in response to 10µM H_2_O_2_ (at the gray line, n>100 cells). **B** Traces of Auranofin (Trx/Grx inhibitor) -induced changes on Grx1-roGFP2 and oROS-HT sensors co-expressed in HEK29. Grx1-roGFP2 responses (green) are shown as the ratio of 510nm emission at 405 and 488nm excitation. oROS-HT’s responses (magenta) are shown as relative fluorescence change from the baseline (ΔF/Fo). Auranofin or vehicle was applied shortly after the start of the imaging sessions (gray dashed line). The dotted trace for each sensor plot represents responses to vehicle treatment (n>100 cells per condition). **C** Translocation of Nrf2 (in nuclear/perinuclear ratio) quantified from immunofluorescence staining of Nrf2 in HEK293 cells exposed to 1µM auranofin for 30 minutes or negative controls. Detailed description of analysis method is in Supp. Fig. 2. **D-I** Dual imaging of Ca^2+^ (Fluo-4, green fluorescence calcium dye) and H_2_O_2_ in hiPSC-CMs in response to 5µM Auranofin. (n=12 for Auranofin, n=11 for Vehicle conditions, ROIs from 3 different biological replicates) **D** Representative traces of oROS-HT (magenta) with Fluo-4 (green) in hiPSC-cardiomyocytes. Peaks (Blue dots) and troughs (red crosses) of Ca^2+^ transients (CaT) are labeled. **E** oROS-HT ΔF/Fo at last frames. **F** Change in resting fluorescence intensity (ΔBaseline) of Fluo-4 from the start to the end of the imaging. **G** Representative CaT changes over time. **H** Schematic description of the CaT phenotypes extracted. **I** Characterization of CaTs from early (first 70 seconds) and late segments(last 70 seconds) of Auranofin and Vehicle treated hiPSC-CM groups: **left** Amplitude (amplitude of CaT at the peak), **middle** TtP (Time-to-Peak), and **right** CaTD90 (CaT Duration 90). **Descriptive Statistics:** Error bars and bands represent the bootstrap confidence interval (95%) of the central tendency of values using the Seaborn (0.11.2) statistical plotting package. **Inferential Statistics:** t-test independent samples. *P < 0.05, **P < 0.01, ***P < 0.001. ****P < 0.0001.

#### Acute effect of auranofin on peroxide and calcium dynamics in hiPSC-CM

There is growing evidence of a mutual interplay between redox and calcium ion dynamics in cells^38^. Ca^2+^ is required in excitable cells such as neurons and cardiomyocytes. Still, simultaneous real-time observations of oxidative stress and Ca^2+^ in the same cell with a temporal resolution that can capture dynamic Ca^2+^ transients (CaT) have been limited. Here, we performed multiplexed imaging of H_2_O_2_ and CaT using oROS-HT_635_ with Fluo-4, a Ca^2+^-sensitive green fluorescent dye in hiPSC-CMs **[Fig. 4D]**. It is widely accepted that oxidative stress perturbs key Ca^2+^ transporters like ryanodine receptors (Ca^2+^ sarcoplasmic reticulum leak)^39^, L-type calcium channels (ICaL, inward Ca^2+^ current)^40^, and sarcoplasmic reticulum calcium ATPase pumps (SERCA, decreased Ca^2+^ reuptake)^41–43^. Functional influence of these perturbations can manifest as changes in specific CaT phenotypes such as baseline Ca^2+^level, CaT amplitude, Time-to-Peak (TtP, on-kinetics), and Calcium Transient Duration 90% (CaTD90, completion of 90% of one CaT period). We explored how the auranofin-induced acute oxidative stress perturbs these transporters and affects Ca^2+^dynamics in detail. Previous studies reported auranofin-induced Ca^2+^ increases in some cell types^44,45^. Indeed, auranofin (5µM) induced peroxide increase **[Fig. 4E]** during the 20-minute imaging period, accompanied by an increase in basal Ca^2+^level **[Fig. 4F]**. Next, we extracted the CaT profile from the Fluo-4 imaging data to further characterize the effect of auranofin **[Fig. 4G, H]**. Compared to the vehicle control, CaTs of auranofin-treated hiPSC-CM exhibited the following phenotypes: elevated CaT peak amplitude and prolonged TtP and CaTD90 **[Fig. 4I]**.

#### Modeling effect of perturbed Ca^2+^ transport on cytosolic Ca^2+^ levels in silico

To investigate whether the oxidative insult and their effects on calcium transporters would lead to the observed changes in the CaT phenotypes, we simulated the intracellular calcium level dynamics using a pre-existing model for CaT in iPSC-CMs^46^. Aligned with the reported effect of oxidative stress on the calcium transporters discussed above, we modified parameters corresponding to the cytosolic calcium efflux via SERCA, the SR Leak amplitude, and the conductance of the L-type Ca^2+^ channel (ICaL) to model oxidative stress. The trend in simulation aligned with observed CaT phenotypes: decreased SERCA uptake simulated a pronounced increase in intracellular calcium baseline, delay of TtP and CaTD90, while higher ICaL conductance showed a pronounced increase in intracellular calcium baseline and CaT amplitude. Interestingly, increased SR Leak did not noticeably affect the aforementioned CaT phenotypes **[Supp. Fig. 3]**, reflecting the hiPSC-CMs electrophysiological immaturity. Specifically, CaTs in hiPSC-CM models are often mostly governed by L-type calcium channel activities due to functional immaturity associated with SR-associated calcium transporters^47–51^, which may explain the observed CaT insensitivity to the increased SR leak. This result is further supported by our 11×10 synergistic perturbation simulation of ICaL and SERCA **[Supp. Fig. 4A].** The baseline calcium level showed pronounced elevation with a focal point at SERCA 0.5x and ICaL 2.0x activity levels (relative to the starting conditions). In contrast, the CaT amplitude showed an elevated focal point around SERCA 0.25x, ICaL 2.0x activity levels. The focal point for TtP and CaTD90 elevation lies near SERCA 0.1x, ICaL 1.0x activity levels. We calculated a CaT influence map derived from an additive weighing of the normalized individual phenotype arrays. It revealed the biased influence of ICaL over SERCA for the phenotypic changes, implying that observed CaT phenotypes in the study may be the result of the biased effect of ICaL over SERCA for calcium handling **[Supp. Fig. 4B].** The result acknowledges the potential intricate nature of effect of oxidative stress on calcium dynamics in cardiomyocytes, which calls for systemic studies on the influence of oxidative stress on specific calcium transport and their synergistic outcome.

### Multiparametric imaging of intracellular and extracellular peroxide dynamics

oROS-HT_635_ could be targeted to other cellular subcompartments, including the mitochondrial matrix, mitochondrial intermembrane space, actin cytoskeleton, and intracellular side of the plasma membrane, using trafficking sequences **[Fig. 5A]**. Intracellular H_2_O_2_ generation is potentially localized and functionally differentiated in aerobic organisms^52^, which calls for monitoring of H_2_O_2_ in a spatially resolved manner (e.g. cellular sub-compartments)^15^. Growing evidence demonstrates the significant contribution of NADPH oxidase-sourced superoxide and peroxide in redox signaling and disease progression^53–57^. The oxidase generates H_2_O_2_ on the extracellular side of the cellular plasma membrane^58^, constituting an extracellular pool of H_2_O_2_ ^59^. Its distribution is achieved through autocrine (aquaporin-mediated diffusion of peroxide^60,61^ into cells) and paracrine^62^ mechanisms. oROS-HT fused to PDGFR transmembrane domain-based trafficking sequence (pDisplay vector, invitrogen) showed robust membrane localization of oROS-HT, and its co-expression with oROS-G, a green variant of oROS-HT we previously reported^25^, was well tolerated in HEK293 **[Fig. 5B]**. Here, we measured menadione-induced H_2_O_2_ increase in both extracellular and intracellular space. Intriguingly, we found that the extracellular peroxide response detected by oROS-HT (inverse response sensor) was faster than oROS-G (direct response sensor). This supports previous observations that menadione increases H_2_O_2_ in the extracellular space, potentially via NADPH oxidase-sourced peroxide^63–66^ **[Fig. 5C]**.

**Figure 5.**
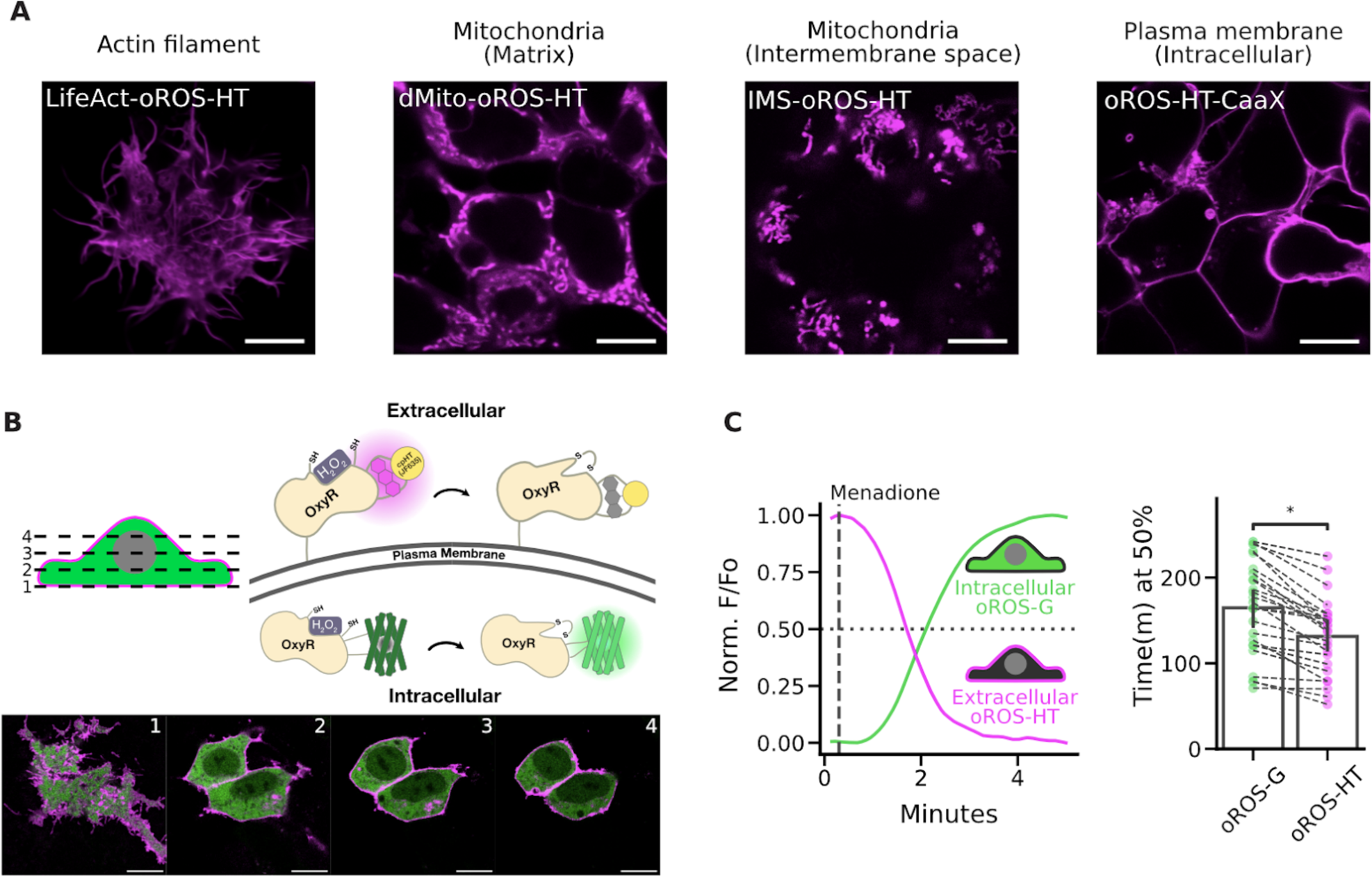
Multi-parametric, dual-color imaging of intracellular and extracellular peroxide dynamics. **A** Subcellular localization of oROS-HT was achieved by previously reported trafficking sequences for actin (LifeAct), mitochondrial matrix (dMito), mitochondrial intermembrane space (IMS), and intracellular side of the plasma membrane (CaaX). HEK293 expressing each of the variants were live-imaged using a Leica SP8 confocal microscope. Scale bar: 10µM. **B** Confocal z-stack images of HEK293 cells co-expressing pDisplay-oROS-HT (extracellular side of the plasma membrane) and pC1-oROS-G (intracellular). Scale bar: 10µM. **C** Fluorescence change of HEK293 expressing pDisplay-oROS-HT and pC1-oROS-G in response to 25 µM Menadione imaged with epifluorescence microscope. Both sensors were imaged every second. ***Left*** A representative trace of oROS-HT and oROS-G from a single cell. ***Right*** time (in minutes) at 50% sensor activation (n=25 cells from 4 biological replicates). **Inferential Statistics:** t-test independent samples. *P < 0.05, **P < 0.01, ***P < 0.001. ****P < 0.0001.

## Discussion

This study introduces a novel bright far-red chemigenetic indicator for peroxide, oROS-HT_635_. To fully harness the brightness of JF635 rhodamine dye, this inverted response sensor was further optimized for higher brightness and dynamic range while exhibiting unmatched sensitivity and kinetics compared to existing red-shifted GEHIs. Since oROS-HT_635_ maintains bright fluorescence in the sensor activation range (e.g. partially oxidized state), it detects high-fidelity signal at physiological peroxide levels. By incorporating HaloTag labeled with JF635 as its reporter, we could achieve oxygen-insensitive, pH-resistant, and photochromic artifact-free imaging that vastly extends its application range. Guided by the crystal structures of OxyR, we optimized the peroxide sensing efficiency of oROS-HT_635_, implying the design avoids disruption of the flexible protein region critical for H_2_O_2_-induced disulfide bridging.

Harnessing oROS-HT_635_’s exceptional multiplexing capability, we performed imaging paired with green fluorescence-based redox potential and Ca^2+^ sensors, allowing monitoring of peroxide level, along with changes in redox potential or calcium. Auranofin, a treatment for rheumatoid arthritis, is gaining attention from the cancer community as a potential therapeutic candidate due to its dose-and-cell-dependent multifaceted mode of action^67,68^. As a Trx/Grx inhibitor, it attenuates the intracellular antioxidant capacity, which increases oxidative stress. Intriguingly, recent studies to repurpose auranofin as a potential cancer therapeutics revealed a more nuanced role of auranofin as increasing cellular oxidative stress can activate regulators such as Nrf2 to boost cellular antioxidative capacity^67,68^. Here, we showed, in real-time, how low-dose auranofin initiates transient oxidative stress, followed by a Grx-independent reversal of H_2_O_2_ levels. The time course of the reversal correlated with increased Nrf2 translocation into the cell nucleus in HEK293 cells, supporting observations from previous studies. Auranofin also altered dynamic calcium transients (CaT) in hiPSC-cardiomyocytes (hiPSC-CM), correlating with an increased level of H_2_O_2_. These observations were consistent with our computational simulation of the effect of oxidative stress on key Ca^2+^ transporters. They confirmed previous studies identifying tight coupling between oxidative stress and calcium transport in various cells and tissues^38,39,43^.

Users can also exploit the remarkable subcellular targeting of oROS-HT_635_ to monitor peroxide with higher spatial resolution near its sources. GEHIs have been pivotal in unraveling cellular peroxide topology by enabling optical monitoring of peroxide dynamics in spatially resolved manner in cytoplasmic and mitochondrial spaces^11,69^. Users could study peroxide biology by disintegrating the topology of peroxide from mitochondria, plasma membrane spaces, and paracrine peroxide^62^, which is critical for understanding the systemic propagation of peroxide build-up in tissues and organisms. Specifically, membrane-tagged oROS-HT_635_ provides new opportunities to investigate peroxide topology proximal to the plasma membrane, which is well demonstrated by the result that re-highlights the potential involvement of plasma membrane NADPH oxidases in menadione-induced peroxide production^63–66^.

The next iteration of oROS-HT_635_ could be optimized for other JF dyes with shifted emission spectra ranging from (494 nm to 722 nm), further enhancing its flexibility in multiplexed optogenetic applications. Another possible avenue for future oROS-HT_635_ development is maximizing its *in vivo* application capability. As a trade-off to its exceptional fluorogenicity, the bioavailability of JF635 dye can be a challenge for animal application. We envision two paths for optimizing the use of oROS-HT_635_ in live animals. First, introducing the dye into brain tissue can be aided with engineered solutions such as injection cannulas or drug delivery systems^70,71^. Alternatively, optimization of the oROS-HT_635_ with highly bioavailable dyes (e.g. JF669)^72,73^ can be explored for efficient animal applications.

In conclusion, oROS-HT_635_ enables the monitoring of peroxides with high spatiotemporal resolution, offering unparalleled flexibility in its multiplexed application with other optogenetic tools. The rapid kinetics and robust subcellular targeting capabilities of oROS-HT_635_, particularly at the outer and inner surfaces of the plasma membrane, render it an invaluable tool for investigating peroxide topology near the plasma membrane. When used with fluorescent sensors for various analytes, oROS-HT_635_ facilitates a dynamic, multidimensional analysis of peroxide changes and environmental responses in real-time, enhancing the contextual understanding of peroxides in biological systems.

## Acknowledgments

J.D.L was supported by 1F31DA056121-01A1 and ISCRM Fellowship. A.B was supported by the Brain Research Foundation, UW Royalty Research Fund, UW ISCRM IPA, NIGMS R01 GM139850-01, P30 DA048736-01-Pilot, NIMH RF1MH130391, NINDS U01NS128537, NIDA R21DA051193 and the McKnight Foundation’s Technologies in Neuroscience Award. S.J.W. was supported by the National Science Foundation DGE-2140004 and the Herbold Foundation. K. E was supported by T32AG066574. A.M.A. Was supported by the National Institute of General Medical Sciences grant RM1 GM131981, the National Institute of Arthritis and Musculoskeletal and Skin Diseases grant P30 AR074990, American Heart Association supplement grant AHA872208 and BCTP-NIH – NIBIB - 5T32EB032787-02. We would like to thank the Janelia Materials program from Howard Hughes Medical Institute Janelia research campus for generous sharing of their Janelia Fluors essential for this study. The research received additional support from the Lynn and Mike Garvey Imaging Core, the UW NAPE Center, and ISCRM Shared Equipment. We want to thank Dr. Randy Moon for his support.

## Code Availability

The source code will be available at https://github.com/BerndtLab/oROS-HT_manuscript.

## Ethics Statement

This study was performed in strict accordance with the recommendations in the Guide for the Care and Use of Laboratory Animals of the National Institutes of Health. All animals were handled according to the approved Institutional Animal Care and Use Committee (IACUC) protocols #4422-01, #4383-02 of the University of Washington and followed the National Institute of Health and the 25 Association for Assessment and Accreditation of Laboratory Animal Care International guidelines. The University of Washington has an approved Animal Welfare Assurance (#A3464) on file with the National Institute of Health Office of Laboratory Animal Welfare (OLAW), is registered with the United States Department of Agriculture (USDA, certificate #91-R-0001), and is accredited by American Association for Accreditation of Laboratory Animal Care International.

## Material requests

Plasmids for oROS-HT and its loss-of-function (C199S) and subcellular targeting variants described in this paper will be available through Addgene: pC1-lifeact-oROS-HT (#216420), pC1-IMS-oROS-HT (#216419), pC1-dmito-oROS-HT (#216418), pC1-oROS-HT-CaaX (#216417), pDisplay-oROS-HT (#216416), AAV2_CAG_oROS-HT(C199S)_WPRE (#216415), AAV2_CAG_oROS-HT_WPRE (#216414), pCAG_oROS-HT-WPRE (#216413), pCAG_oROS-HT_LF(C199S)-WPRE (#216412). Authors will also provide plasmids upon request.

## Methods

### Molecular Biology

oROS-HT variants were all cloned based on the pC1 plasmid backbone from pC1-HyPer-Red (Addgene ID: 48249). Primers for point mutations or fragment assembly required to generate the oROS-HT screening variants were designed for In Vitro Assembly cloning (IVA) technique^74^, and they were ordered from Integrated DNA Technologies (IDT). All gene fragment amplifications were done using Superfi-II polymerase (Invitrogen; 12368010). Amplification of the DNA fragment was verified with agarose gel electrophoresis. 30 minutes of DpnI enzyme treatment were done on every PCR product to remove the plasmid template from PCR samples. Circulaization or assembly of the PCR products was achieved with the IVA technique, while the linear DNA products were transformed into competent E.Coli cells (DH5ɑ or TOP10) and grown on agar plates that contain kanamycin selection antibiotic (50 µg/mL). Upon colony formation, single colonies were picked and grown in 5mL cultures containing LB Broth (Fisher BioReagents; BP9723-2) and selection antibiotic (/kanamycin; 50 µg/mL) overnight (37°C, 230 RPM). DNA was isolated using Machery Nagel DNA prep kits (Machery Nagel; 740490.250). Sanger sequencing (Genewiz; Seattle, WA) or Whole-plasmid nanopore sequencing (Plasmidsarus; Eugene, OR) of the isolated plasmid DNA was used to confirm the presence of the intended mutation. Genes encoding the final variants were cloned into a CAG-driven backbone, pCAG-Archon1-KGC-EGFP-ER2-WPRE (Addgene; #108423), using the methods above. All subsequences were verified with Sanger sequencing (Genewiz; Seattle, WA) or Whole-plasmid nanopore sequencing (Plasmidsarus; Eugene, OR).

### Protein structure prediction and analysis

Protein structure analysis and plotting were performed using Chimera-X-1.2.1. Oxidized [PDB:1I6A] and reduced [PDB:1I69] crystal structures of ecOxyR were imported from the Protein Data Bank (PDB). Pairwise residue distance between reduced and oxidized ecOxyR structure was achieved by aligning both structures using a matchmaker algorithm that superimposes protein structures by creating a pairwise sequence alignment and then fitting the aligned residue pairs to derive pairwise residue distances. The structure of Variant 213-214 was predicted using ColabFold^19^. (msa_method=mmseqs2, homooligomer=2, pair_msa=False, max_msa=512:1024, subsample_msa=True, num_relax=None, use_turbo=True, use_ptm=True, rank_by=pLDDT, num_models=3, num_samples=1, num_ensemble=1, max_recycles=24, tol=0, is_training=False, use_templates=False). The putative position of JF635 was incorporated into the ColabFold prediction of Variant 213-214 to report JF635 bound cpHaloTag structure (PDB:6U2M) with the matchmaker algorithm.

### Chemicals

Halotag ligand of Janelia Fluor (JF-HTLs) 635, 585 described in this paper were generously provided by Janelia Materials. Stock solutions of JF-HTLs were prepared in 100% DMSO at 200µM. Cells described in this study were incubated in 200nM JF-HTL for 1 hour prior to imaging unless specified. H_2_O_2_ working solutions were freshly prepared before every experiment from H_2_O_2_ solution 30 % (w/w) in H_2_O (Sigma-Aldrich, H1009). A stock solution of Menadione (Sigma-Aldrich, M9429) was prepared in 100% DMSO at 50mM. A stock solution of Auranofin (Tocris Bioscience, 46-005-0) was prepared in 100% DMSO at 50mM.

### HEK Cell culture and transfection

Human Embryonic Kidney (HEK293; ATCC Ref: CRL-1573) cells were cultured in Dulbecco’s Modified Eagle Medium + GlutaMAX (Gibco; 10569-010) supplemented with 10% fetal bovine serum (Biowest; S1620). When cultures reached 85% confluency, the cultures were seeded at 150,000/75,000 cells per well in 24/48-well plates, respectively. 24 hours after cell seeding, the cells were transfected using Lipofectamine3000 (Invitrogen; L3000015) at 1000/500 ng of DNA per well of a 24/48-well plate, according to the manufacturer’s instructions.

### Primary rat neuron isolation

Primary cortical neurons were prepared as previously described^75,76^. Briefly, 24-well tissue culture plates were coated with matrigel (mixed 1:20 in cold-PBS, Corning; 356231) solution and incubated at 4°C overnight before use. Sterile dissection tools were used to isolate cortical brain tissue from P0 rat pups (male and female). Tissue was minced until 1mm pieces remained, then lysed in equilibrated (37°C, 5% CO2) enzyme (20 U/mL Papain (Worthington Biochemical Corp; LK003176) in 5mL of EBSS (Sigma; E3024)) solution for 30 minutes at 37°C, 5% CO2 humidified incubator. Lysed cells were centrifuged at 200xg for 5 minutes at room temperature, and the supernatant was removed before cells were resuspended in 3 mLs of EBSS (Sigma; E3024). Cells were triturated 24x with a pulled Pasteur pipette in EBSS until homogenous. EBSS was added until the sample volume reached 10 mLs before spinning at 0.7 rcf for 5 minutes at room temperature. The supernatant was removed, and enzymatic dissociation was stopped by resuspending cells in 5 mLs EBSS (Sigma; E3024) + final concentration of 10 mM HEPES Buffer (Fisher; BP299-100) + trypsin inhibitor soybean (1 mg/ml in EBSS at a final concentration of 0.2%; Sigma, T9253) + 60 µl of fetal bovine serum (Biowest; S1620) + 30 µl 100 U/mL DNase1 (Sigma;11284932001). Cells were washed 2x by spinning at 0.7 rcf for 5 minutes at room temperature and removing supernatant + resuspending in 10 mLs of Neuronal Basal Media (Invitrogen; 10888022) supplemented with B27 (Invitrogen; 17504044) and glutamine (Invitrogen; 35050061) (NBA++). After final wash spin and supernatant removal, cells were resuspended in 10 mLs of NBA++ before counting. Just before neurons were plated, matrigel was aspirated from the wells. Neurons were plated on the prepared culture plates at the desired seeding density. Twenty-four hours after plating, 1µM AraC (Sigma; C6645) was added to the NBA++ growth media to prevent the growth of glial cells.Plates were incubated at 37°C and 5% CO2 and maintained by exchanging half of the media volume for each well with fresh, warmed Neuronal Basal Media (Invitrogen; 10888022) supplemented with B27 (Invitrogen; 17504044) and glutamine (Invitrogen; 35050061) every three days.

### Brain slice imaging

#### Organotypic whole hemisphere (OWH) rat brain slice preparation

Male rats on postnatal day (P)10 were administered an overdose intraperitoneal injection of pentobarbital (120–150 mg/kg). Animals were then quickly decapitated and whole brains were extracted, cut into hemispheres, and placed into ice-cold dissecting media consisting of 0.64% w/v glucose, 100% Hank’s Balanced Salt Solution (HBSS), 1% penicillin–streptomycin. Whole-hemisphere live slices of 300 μm were obtained using a tissue chopper as previously described.^77^ Slices were then transferred to 35 mm, 0.4 μm-pore membrane inserts in six-well plates and cultured in 1 ml of 5% heat-inactivated horse-serum slice culture media (SCM) consisting of 50% Minimum Essential Media (MEM), 45% HBSS, 1% GlutaMAX, and 1% penicillin–streptomycin. Slices were cultured in a sterile incubator at constant temperature (37°C), humidity, and CO_2_ level (5%).

#### AAV transduction and confocal imaging

After 1 day in vitro (DIV), crude AAV9-CAG-oROS-HT prep was added to the slices to be expressed. At the end of the 3-day incubation, 1 μM JF635-HTL was added to the slices for an additional 48 hours. OWH brain slices were transferred to 35mm confocal dishes (VWR, 75856-742). Confocal images were acquired with 10x (Nikon Plan Apo 10x Objective, 0.45 numerical aperture) and 20x (Nikon Plan Apo 10x Objective, 0.75 numerical aperture) magnifications (Nikon Corporation, Minato City, Tokyo, Japan). Brain slice tile scans were obtained with the Cy5 channel before multiple representative images were taken from both the cortex and striatum of each slice. Image acquisition settings were kept consistent before and after the 300µM H_2_O_2_ stimulation.

### Differentiation of stem cell-derived cardiomyocytes and neurons

#### hiPSC culture and cardiomyocyte differentiation (diffusion study)

Undifferentiated IMR90 (WiCell) hiPSCs were maintained on Matrigel (Corning) coated tissue culture plates in mTeSR1 (Stemcell Technologies). Cardiomyocyte-directed differentiation was performed using a modified small molecule Wnt-modulating protocol using Chiron 99021 and IWP-4 as previously described.^78,79^. Lactate enrichment was performed following differentiation to purify hiPSC-CMs.^80^

#### hiPSC culture and cardiomyocyte differentiation (Auranofin study)

Undifferentiated human induced pluripotent stem cells (hiPSCs) (WTC11, Male) were maintained on Matrigel (Corning) coated tissue culture plates in mTeSR1 (Stemcell Technologies). Cardiomyocyte-directed differentiation was performed using the RBA-based modified method as previously described^81^. Spontaneous contraction was observed on day 8 post-induction. On day 12 post-induction, media was reduced to 1 mL in preparation for 45 minutes heat-shock at 42°C on day 13. After heat shock, the media was changed to 1 mL of fresh RMPI+B27+ins. On day 14, cells were dissociated with 0.05% Trypsin (Thermo-Fisher) and frozen in BAMBANKER for storage in LN_2_. These cardiomyocytes were thawed in 90% RPMI+B27+ins and 10% Knockout Serum (KSR) with 10μM ROCK inhibitor and plated on matrigel coated plates. 24 hours after thaw, media was replaced with fresh RPMI+B27+ins, and changed every other day.

### hiPSC culture and cortical neuron differentiation

Neurons were generated from the previously characterized wild type CV background human induced pluripotent stem cell line^82–84^. Neural progenitor cells (NPCs) from this cell line were differentiated from hiPSCs using dual-SMAD inhibition and NPCs were differentiated into neurons as previously described (Knupp et al., 2020; Shin et al., 2023). Briefly, for cortical neuron differentiation from NPCs, NPCs were expanded into 10 cm plates in Basal Neural Maintenance Media (BNMM) (1:1 DMEM/F12 (#11039047 Life Technologies) + glutamine media/neurobasal media (#21103049, GIBCO), 0.5% N2 supplement (# 17502-048; Thermo Fisher Scientific,) 1% B27 supplement (# 17504-044; Thermo Fisher Scientific), 0.5% GlutaMax (# 35050061; Thermo Fisher Scientific), 0.5% insulin-transferrin-selenium (#41400045; Thermo Fisher Scientific), 0.5% NEAA (# 11140050; Thermo Fisher Scientific), 0.2% β-mercaptoethanol (#21985023, Life Technologies) + 20 ng/mL FGF (R&D Systems, Minneapolis, MN). Once the NPCs reached 100% confluence, they were switched to Neural Differentiation Media (BNMM +0.2 mg/mL brain-derived neurotrophic factor (CC# 450–02; PeproTech) + 0.2 mg/mL glial-cell-derived neurotrophic factor (CC# 450–10; PeproTech) + 0.5 M dbcAMP (CC# D0260; Sigma Aldrich). Neural Differentiation Media was changed twice a week for 21 days, at which point the differentiation is considered finished. Neurons were replated at a density of 500,000/cm^2^.

### Immunofluorescence staining

Immunofluorescence staining performed for Nrf2 translocation study were done using polyclonal Nrf2 antibody (PA5-27882, Invitrogen) and Donkey anti-Rabbit IgG Alexa Fluor 488 (A21206, Invitrogen). HEK293 cells for each condition were fixed in 4% paraformaldehyde for 15 minutes and permeabilized in 0.2% Triton-x solution for 1 hour. After blocking the fixed cells for 1 hour with 0.5% Bovine Serum Albumin (BSA) blocking buffer in TBST, Cells were then incubated with primary antibodies diluted in the blocking buffer overnight at 4°C. The next day, cells were washed 3 times with PBS. They were then incubated in a secondary antibody solution containing secondary antibodies diluted in 0.5% BSA in PBS overnight at 4°C. Counterstaining was performed with Vectashield containing DAPI (Vector Labs).

### Microscopy

Imaging experiments described in this study were performed as follows unless specifically noted. Epifluorescence imaging experiments were performed on a Leica DMI8 microscope (Semrock bandpass filter: GFPratio ex/em: FF01-391-23/FF01-520-35, GFP ex/em: FF01-474-27/FF01-520-35, RFP ex/em:FF01-554-23 or FF01-578-21/FF01-600-37, Far-red ex/em: FF01-635-18/FF01-680-42) controlled by MetaMorph Imaging software, using sCMOS camera (Photometrics Prime95B) and 20x magnification lens (Leica HCX PL FLUOTAR L 20x/0.40 NA CORR) or 10× objective (Leica HC PL FLUOTAR L 10x/0.32 NA) Confocal imaging experiments were performed on Leica SP8 confocal microscope from Imaging Core of Institute of Stem Cell and Regenerative Medicine. Cells were imaged in live cell imaging solution with 10mM glucose (LCIS+, Gibco, A14291DJ). Image analysis methods are described below.

### Hypoxic oROS-HT sensor maturation in HEK293

2 day post-seeding of HEK293 cells in 24 well plates (150,000 cells/well), culture media was swapped from complete DMEM media (as mentioned above) to complete Fluorobrite DMEM (A1896701, Gibco) with 20mM HEPES. After 2-hour of acclimation, cells were transfected (Lipofectamine-based, as described above) with pC1-oROS-HT-C199S (Loss-of-function), with 100nM JF635-HTL. Immediately after the transfection, transfected cells were either incubated at 37°C in an atmospheric environment or under hypoxic conditions. For hypoxic conditions, culture plates were transferred into a sealable chamber. The chamber was flushed with N_2_ for 10 min at a flow rate of 10 L/min before being placed into the incubator. Approximately 18 hours after, epifluorescence imaging were performed as described earlier.

### Multiplexed experiments

#### oROS-HT/SypHer3s

HEK293 cells were co-transfected with pC1-oROS-HT/pC1-SypHer3s or pC1-oROS-HT-C199S/pC1-SypHer3s as described above. 2 days post-transfection, both sensors expressed in HEK293s were imaged using epifluorescence microscope. pH change experiment for oROS-HT-C199S were performed with HEK293s in PBS (10010001, Gibco) prepared at pH of 6, 7.44, and 9. Fluorescence level for GFP and Far-red profile were captured every 1.5 seconds. Sequential pH-changes plus Menadione applications were performed with HEK293s in PBS (pH 7.44), which was changed to PBS (pH 6) followed by menadione stimulation prepared in PBS (pH 6). Fluorescence level for GFP and Far-red profile were captured every 2 seconds.

#### oROS-HT/Grx1-roGFP2

HEK293 cells were co-transfected with pC1-oROS-HT and pC1-Grx1-roGFP2 as described above. 2 days post-transfection, both sensors expressed in the cells with live cell imaging solution with 10mM glucose (LCIS+, Gibco, A14291DJ) were imaged using an epifluorescence microscope. For the sequential response of oROS-HT/Grx1-roGFP2 to 10µM H_2_O_2_, fluorescence level for GFP and Far-red profile were captured every second. For the response to Auranofin, fluorescence levels for GFPratio, GFP, and Far-red profiles were captured every minute.

#### oROS-HT/Fluo-4

hiPSC-CMs were transfected with pCAG-oROS-HT as described above. 2 days post-transfection, cells were incubated with Fluo-4 (Invitrogen, F14201) at 5µM and JF635-HTL in RPMI + B27+insulin for 1 hour prior to imaging. For the response to Auranofin, fluorescence level of GFP profile (10Hz) and Far-red (0.1Hz) profile were acquired every 10 seconds for hiPSC-CMs in HEPES-buffered RPMI + B27+insulin.

### Analysis

Analysis of cell fluorescence imaging data was done by FUSE, a custom cloud-based semi-automated time series fluorescence data analysis platform written in Python. First, the cell segmentation quality of the selected Cellpose^85^ model was manually verified. For the segmentation of cells expressing cytosolic fluorescent indicators, model ‘cyto’ was selected as our base model. If the selected Cellpose model was low-performing, we further trained the Cellpose model using the Cellpose 2.0 human-in-the-loop system^86^. Using an “optimized” segmentation model, fluorescence time-series data is extracted for each region of interest. This allows for unbiased extraction of change in cellular fluorescence information for a complete set of experimental samples. Extracted fluorescence data is normalized as specified in the text using a custom Python script.

### Computational Cell Scale Modeling

We used an existing model of iPSC-CM membrane kinetics^46^ with one modification. Based on experimental observations, the spontaneous beating of the iPSC-CMs was observed to be around 0.5 Hz. To reflect this observation in our computational simulations, we increased the maximal value of the inward rectifier potassium (I_K1_) by a factor of 1.71484375. This change resulted in a decrease in spontaneous beating rate from 1.1 Hz to 0.5 Hz. To simulate ROS effects on iPSC-CMs, we ran simulations in which we modified parameters corresponding to maximal efflux via the SR Ca^2+^ ATPase (SERCA2a), SR Leak amplitude, and maximal conductance of the L-type Ca^2+^ channel (g_CaL_). The perturbation factor for SERCA efflux varied from 0.1 to 1.0 in steps of 0.1. SR Leak amplitude and g_CaL_ were both increased from the default level (1.0) to 2.0 in steps of 0.1. Simulations of bioelectrical activity were conducted using openCARP^87^, a cardiac electrophysiology modeling software that is freely available for non-commercial reuse (see: http://opencarp.org/). Stimulated Ca_i_ values were post-analyzed with custom-written python scripts. Scripts and files used to run all simulations can be found at aforementioned Github depository.

