## Supplemental Files for "Far-red and sensitive sensor for monitoring real time H_2_O_2_ dynamics with subcellular resolution and in multi-parametric imaging applications"

**Supplementary Figure 1** Alphafold-guided identification of mutational hotspot of oROS-HT.

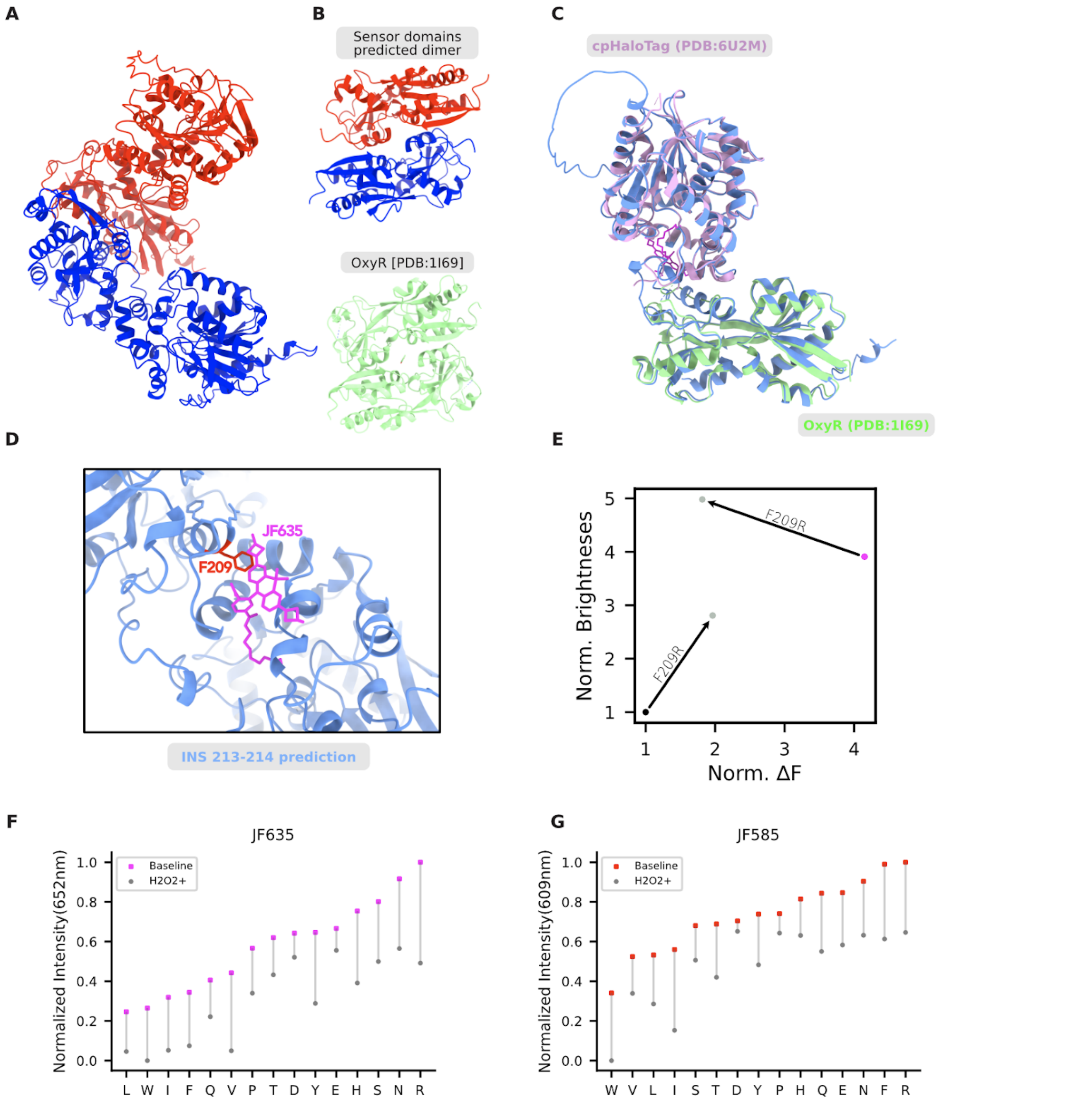


**A** Structure prediction of cpHaloTag 213-214 insertion variant in a dimeric state using ColabFold (AlphaFold + MMseqs2). **B** ***top*** Sensing domains (OxyR) of each dimeric pair from cpHaloTag 213-214 insertion variant ColabFold prediction. ***bottom*** Experimentally resolved crystal structure of reduced OxyR in a dimeric state [PDB:1I69]. **C** Superimposed structures of cpHaloTag 213-214 insertion variant and cpHaloTag with JF635 ligand (reporting domain, extracted from PDB: 6U2M) and OxyR (sensing domain, extracted from PDB: 1I69). **D** Proximity of residue F209 to JF635 fluorophore of superimposed cpHaloTag structure. **E** Based on Alphafold2 predicted structure of cpHaloTag 213-214 insertion variant, a mutational target of F209 was identified based on proximity. F131R mutation yielded in higher fluorescence baseline. (black dot). Consitantly, F209R yielded higher baseline fluorescence for oROS-HT (magenta dot) but it significantly diminished the dynamic range of the sensor. **F** Mutational screening of residue 209 in cpHaloTag 213-214 insertion variant with JF635 in HEK293 WT cells. (n>100 cells per variant). **G** Mutational screening of residue 209 in cpHaloTag 213-214 insertion variant with JF585 in HEK293 WT cells. (n>100 cells per variant).

**Supplementary Figure 2** Nrf2 translocation quantification.

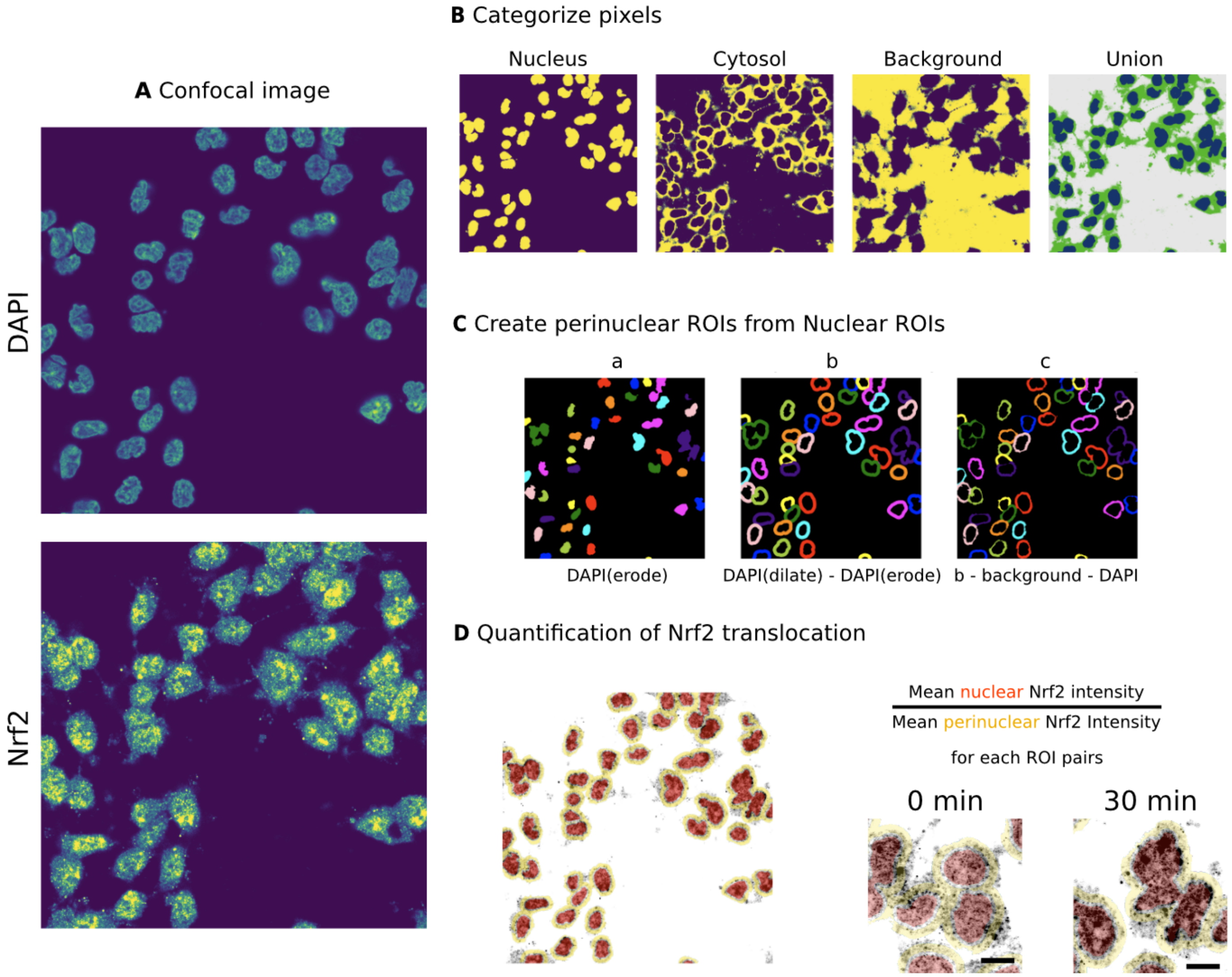


HEK293 cells were fixed (at 0 or 30 minutes upon Auranofin 1μM exposure), then an immunofluorescence assay was performed for Nrf2 (see methods section). **A** Next, fixed cells were imaged under a confocal microscope for Nrf2 subcellular localization. Post-processing of the images to quantify Nrf2 translocation was done using a custom analysis method. **B** We sorted each small part of the image (pixel) into 3 groups. If pixel is positive for DAPI, it is considered "Nucleus". If pixel is negative for DAPI but positive for Nrf2, it is considered "Cytosol". Pixels that are not nucleus nor cysosol are considered background. **C** We made the region of interest (ROI) that show the nucleus smaller (eroded, a) to precisely measure how much Nrf2 protein is in the nucleus. We expanded (dilated) the nucleus areas to create a ring around them (b). This "perinuclear ring" was further processed to exclude any pixels thats positive for DAPI or background, which helps us measure the amount of Nrf2 protein outside the nucleus, in the cytosol. **D** Lastly, mean nuclear and perinuclear Nrf2 intensities were calculated. The Nrf2 translocation is represented by the change in their ratio for each nuclear and perinuclear ROI pairs (see **Fig. 4C**). Bottom right images represent cells from 0 and 30 minutes upon Auranofin exposure. Scale bar = 10μM.

**Supplmentarty Figure 3** Modeling the effect of perturbed $Ca^{2+}$ transport on $Ca^{2+}$ transients *in silico*.

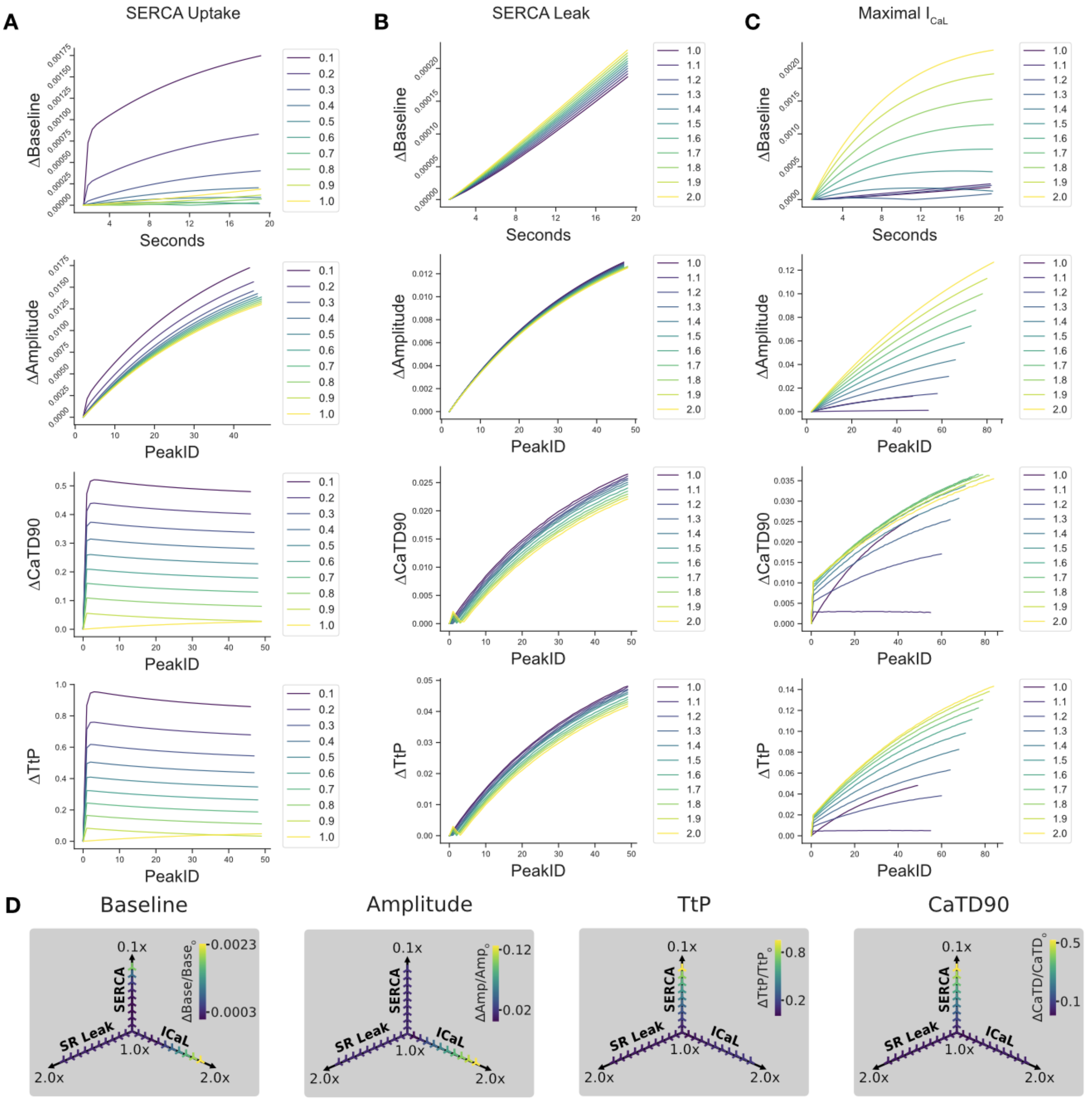


**A-C** *in silico* CaT characterization (Kernik hiPSC-CM model, 0.5Hz initial beating frequency for 20 seconds) in response to graded perturbation of model parameters (color-coded graded changes with respect to the parameter's default value in the Kernik model: 1.0x) for maximal $I_{CaL}$, SERCA uptake, and SR Leak. The following CaT features were extracted from simulation results: ΔBaseline $Ca_i$ level, ΔCaT Amplitude, ΔTtP, and ΔCaTD90. **A** Effect of graded SERCA uptake perturbation on ΔBaseline, ΔAmplitude, ΔTtP, ΔTtB and ΔCaTD90 extracted from the Kernik hiPSC-CMs model simulation. **B** Effect of graded SR leak perturbation on ΔBaseline, ΔAmplitude, ΔTtP, ΔTtB and ΔCaTD90 extracted from Kernik hiPSC-CMs model simulation. **C** Effect of graded CaL conductance perturbation on ΔBaseline, ΔAmplitude, ΔTtP, ΔTtB and ΔCaTD90 extracted from the Kernik hiPSC-CMs model simulation. **D** 3D visualizations of the effects of perturbing each calcium transporter parameter are shown for each feature. Each axis represents graded changes with respect to the parameter's default value in the Kernik model (1.0x).

**Supplementary Figure 4** Modeling the effect of perturbed $Ca^{2+}$ transport on $Ca^{2+}$ transients *in silico*.

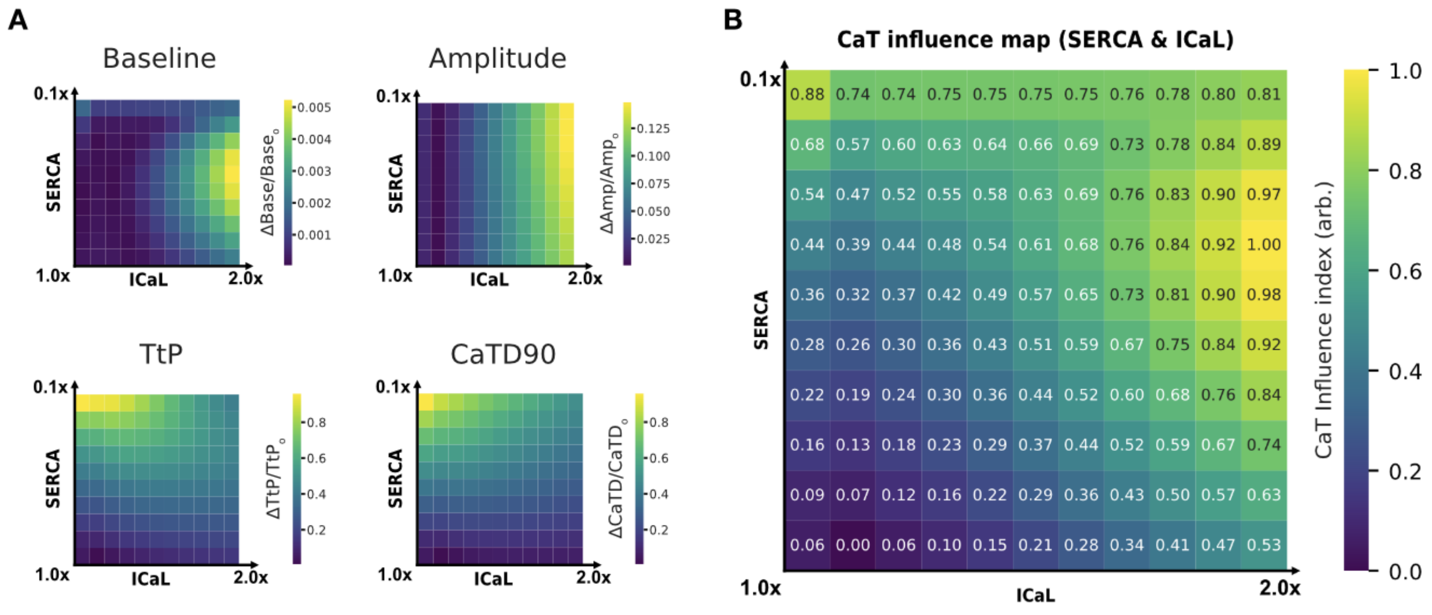


**A** 11x10 (ICaLxSERCA) grid representation of normalized ΔBaseline, ΔAmplitude, ΔTtP, ΔTtB and ΔCaTD90 from simulated combinatorial perturbation of calcium transporters. **B** CaT influence map ($I_{CaL}$ vs SERCA) derived from an additive weighing of the normalized individual phenotype arrays.

**Supplementary Figure 5** Mitochondrial specificity of dMito-oROS-HT

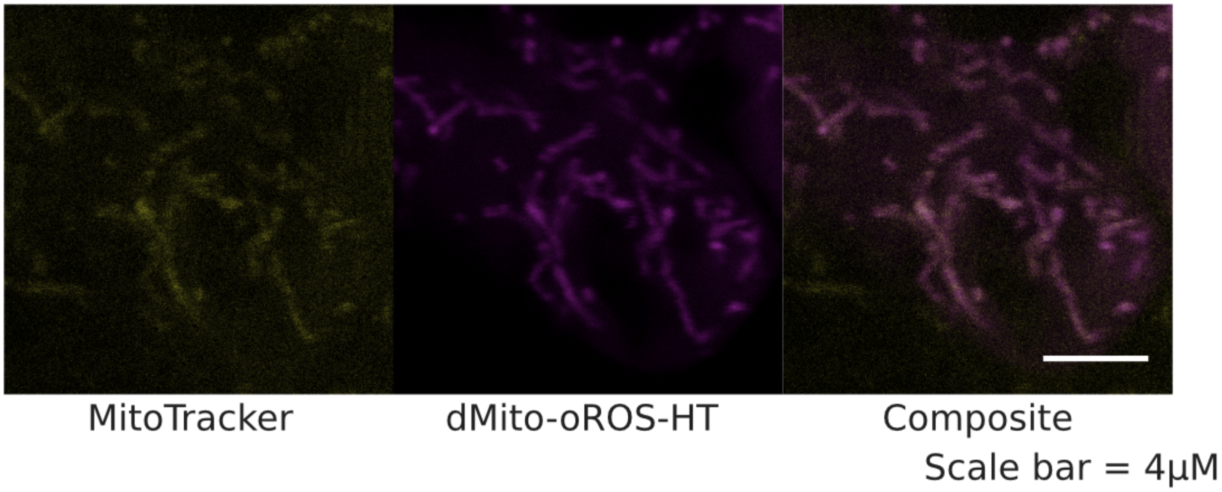


Validation of mitochondrial specificity of dMito-oROS-HT. HEK293 cells expressing dMito-oROS-HT were stained with MitoTracker Orange CMTMRos for a side-by-side comparison of mitochondrial specificity. HEK293 cells expressing each of the variants were imaged using a Leica SP8 confocal microscope.
